# Neural coding of fine-grained object knowledge in perirhinal cortex

**DOI:** 10.1101/194829

**Authors:** Amy Rose Price, Michael F. Bonner, Jonathan E. Peelle, Murray Grossman

## Abstract

**Summary:** Over 40 years of research has examined the role of the ventral visual stream in transforming retinal inputs into high-level representations of object identity [1–6]. However, there remains an ongoing debate over the role of the ventral stream in coding abstract semantic content, which relies on stored knowledge, versus perceptual content that relies only on retinal inputs [7–12]. A major difficulty in adjudicating between these mechanisms is that the semantic similarity of objects is often highly confounded with their perceptual similarity (e.g., animate things are more perceptually similar to other animate things than to inanimate things). To address this problem, we developed a paradigm that exploits the statistical regularities of object colors while perfectly controlling for perceptual shape information, allowing us to dissociate lower-level perceptual features (i.e., color perception) from higher-level semantic knowledge (i.e., color meaning). Using multivoxel-pattern analyses of fMRI data, we observed a striking double dissociation between the processing of color information at a perceptual and at a semantic level along the posterior to anterior axis of the ventral visual pathway. Specifically, we found that the visual association region V4 assigned similar representations to objects with similar colors, regardless of object category. In contrast, perirhinal cortex, at the apex of the ventral visual stream, assigned similar representations to semantically similar objects, even when this was in opposition to their perceptual similarity. These findings suggest that perirhinal cortex untangles the representational space of lower-level perceptual features and organizes visual objects according to their semantic interpretations.

## Highlights

-Perirhinal cortex (PRc) contains combinatorial codes of object category and color

-These PRc codes reflect semantic knowledge of object colors

-V4 encodes color independent of semantic information

-These results reveal a transformation from perceptual to conceptual object codes

## In Brief

Observers of the world know that roses are typically red and violets blue, but neurobiologists know little about how this knowledge is encoded by the brain. Price et al. report a neural mechanism through which humans understand the meaning of object colors, revealing a fundamental biological process that transforms sensory inputs into abstract representations of object meaning.

## Results and Discussion

We designed a novel stimulus set that allowed us to leverage the natural statistics of object-color information in order to investigate the perceptual and semantic coding of visual objects along the ventral stream. Many objects in our natural environment exist in a range of colors and exhibit clear statistical regularities in their color appearance, which have important implications for their meaning (e.g., the color green has a different meaning for leaves than for bananas). The stimuli used in this study were images of objects from three categories (apples, leaves, and roses), which were displayed in five different colors (red, pink, yellow, green, and blue). Example stimuli are shown in Figure 1. The semantic associations of the colors differ in the context of each object category (e.g., green is a common natural color for leaves but not roses). Thus the color-and-object combinations give rise to a unique semantic similarity space for each category, and, importantly, these representational spaces differ from the perceptual similarity space of the colors alone. For example, in the category of apples, green is semantically similar to red (which are both common colors for apples), even though green and red are highly dissimilar in color-perceptual space. We used functional MRI (fMRI) in human participants to measure patterns of neural activity while the subjects viewed images of these objects and performed an unrelated visual-detection task (Supplementary Figure S1).

**Figure 1.**
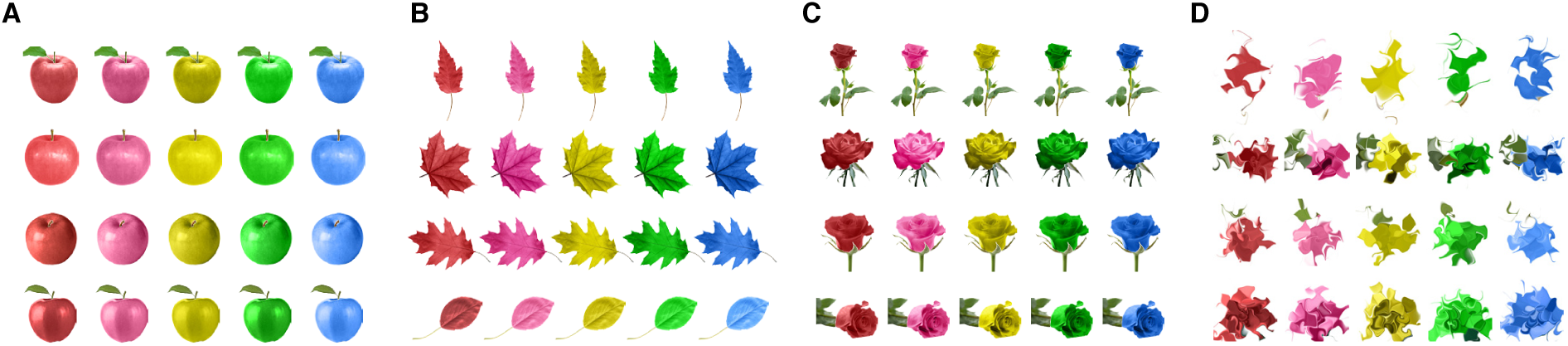
Example stimuli from each object category in each color combination. (A) apples (B) leaves (C) roses (D) non-object diffeomorphically warped images.

We examined two theoretically motivated regions of interest along the ventral stream: V4 and perirhinal cortex (PRc). V4 is a mid-level association region that has previously been implicated in color perception [13–15], but whose possible role in color semantics remains unknown. PRc is a subregion of the anterior temporal lobe located at the apex of the ventral visual pathway [6] that has been implicated in object individuation and object semantics [16–24]. However, it is not yet known whether PRc contains a mechanism for untangling the similarity space of lower-level perceptual inputs and organizing objects according to their semantic interpretations, even when this is at odds with their perceptual similarity.

**Figure 2.**
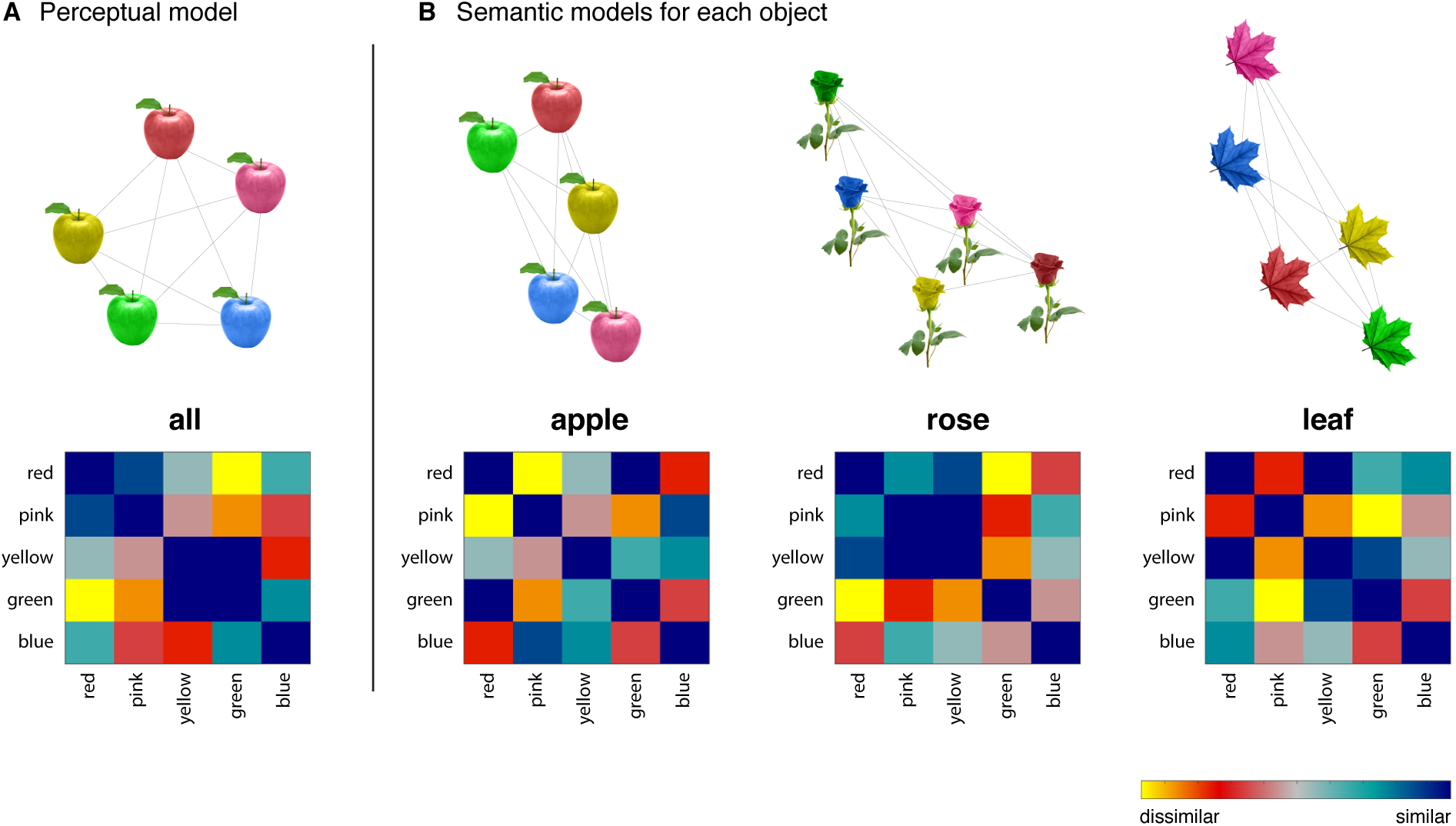
Representational dissimilarity models for the perceptual color model and the category-specific semantic models. Each plot shows a dissimilarity matrix on the bottom and a two-dimensional embedding of stimuli for that model on the top. (A) Perceptual- color model. This model is based on the perceptual similarity of the colors alone and is thus the same for all object categories. The apple category is shown as an example. (B) Semantic color models reflected category-specific co-occurrence statistics and thus are unique for each object category.

We used representational similarity analysis (RSA) to probe the information encoded in the multivariate activity patterns in our regions of interest. We specifically examined the fit of two models that capture the perceptual and semantic similarity of the colors within each category (Figure 2). A key aspect of this design is that we specifically modeled representational dissimilarities within each object category (e.g., roses). This approach perfectly controls for shape information, allowing us to examine the coding of object semantics in a manner that is completely independent of perceptual confounds related to shape.

This analysis revealed a strong double dissociation between the perceptual representation of object colors in V4 and the semantic representation of object colors in PRc (Figure 3; 2 × 2 repeated-measures ANOVA interaction of region by model, F(1,15) = 14.9, p = 0.001). As hypothesized, the semantic similarity model fit significantly in PRc but not V4 (V4: t(15) = 0.87, p = 0.40, Cohen’s d = 0.22; PRc: t(15) = 5.41, p <0.001, Cohen’s d = 1.35), whereas the perceptual color model fit significantly in V4 but not PRc (V4: t(15) = 2.22, p = 0.04, Cohen’s d = 0.56; PRc: t(15) = 0.54, p = 0.60, Cohen’s d = 0.13). Furthermore, direct comparisons showed that the semantic similarity model fit significantly better in PRc than in V4 (t(15) = 3.87, p = 0.002; Cohen’s d = 1.31) and the perceptual similarity model fit significantly better in V4 than in PRc (t(15) = 2.13, p = 0.05; Cohen’s d = 0.67).

**Figure 3.**
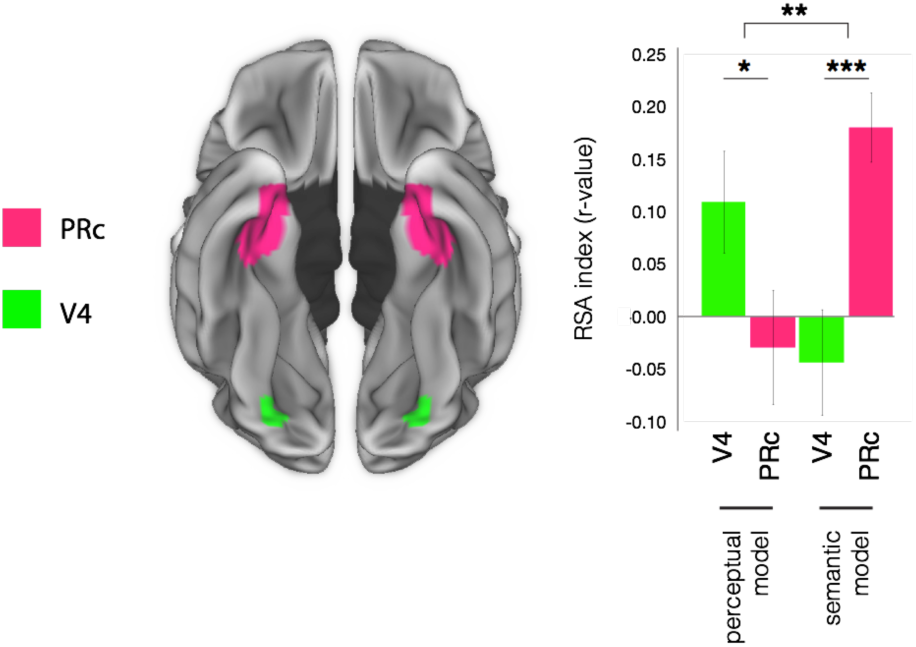
Double dissociation between the processing of color at a perceptual level in V4 and the processing of color at a semantic level in perirhinal cortex. *p < 0.05, **p < 0.01, ***p < 0.001. Plots depict correlation means ±SE.

These findings indicate that PRc integrates the shape and color features of visual objects and links these perceptual inputs with knowledge representations of object colors. In doing so, PRc appears to embed visual object representations in a semantic space that is orthogonal to the lower-level perceptual similarity space of object features. These results are remarkably consistent with a previous neuropsychological report of a patient with a lesion encompassing PRc who had a profound deficit in object-color knowledge but a relative sparing of color perception [25]. These findings are also broadly consistent with previous studies of the semantic variant of primary progressive aphasia (also known as semantic dementia), a neurodegenerative disease that encompasses PRc and results in a profound impairment in object meaning with a relative sparing of visual-perceptual abilities [26, 27]. Furthermore, the results of our color-perceptual model align well with recent work implicating V4 in the coding of a fine- grained perceptual color space [13, 28].

To test for possible effects in other regions of the ventral stream, we performed the same analyses in a series of regions along ventral occipital-temporal cortex (including early visual cortex, lateral occipital complex, inferior temporal gyrus, and fusiform gyrus). We found no significant effects in any other ventral visual regions for either the perceptual color model or the semantic model (Supplementary Figure S2B-C). We also performed an analysis to test for possible univariate effects related to the typicality of color-and-category combinations, which might reflect a coarse familiarity signal in PRc. We found no reliable correlation between color frequency and the mean univariate signal in either PRc or V4 (Supplementary Figure 3S).

Much of the work examining high-level semantic coding in the ventral visual stream has focused on the representation of broad categories of objects (e.g., animate versus inanimate; fruit versus vegetables). However, category membership is often highly correlated with basic shape information [9, 11, 12]. Furthermore, semantic representations of objects encompass much more than their broad category labels. An essential aspect of the semantic memory system is information about individual objects *within* a category. Here we were able examine the coding of fine-grained object semantics in a manner that completely controls for the contribution of perceptual shape information. Interestingly, we did not observe evidence for the coding of fine-grained object semantics in inferior temporal cortex, a region that is strongly associated with the representation of object categories but whose contribution to high-level object semantics has long been debated [7–12]. Rather this more fine-grained semantic information appears to be encoded in PRc—a higher-level region of the ventral stream that has previously been implicated in the detailed analysis that underlies object individuation [17, 19–21, 29, 30]. Our findings are consistent with the proposed role of PRc in object individuation, but they suggest that the mechanism for object representation in PRc is more complex than a perceptual analysis of feature conjunctions. Specifically, these findings suggest that PRc not only disambiguates perceptually confusable objects (e.g., green apple and blue apple) but also assigns similar representations to perceptually distinct objects with similar meanings (e.g., green apple and red apple). Thus, PRc appears to untangle the similarity space of lower-level perceptual inputs and organize individual objects according to their semantic interpretations.

## Author contributions

A.R.P, M.F.B, J.E.P, and M.G. designed the experiment, reviewed the analyses, and discussed the results. A.R.P. and M.F.B. collected the data and performed the analyses. A.R.P, M.F.B, J.E.P, and M.G. wrote the paper.

## Acknowledgments

This work was supported by NIH grants R01 AG017586, the Wyncote Foundation, and the Jameson-Hurvich fund. The authors thank B. Stojanoski for sharing code for diffeomorphic transformations, and R. Epstein for helpful comments on this project.

## Experimental Procedures

### Participants

Sixteen healthy participants (7 female; mean age = 24.6, SD = 2.6) with normal or corrected-to-normal vision were recruited from the University of Pennsylvania community. Participants provided written informed consent in compliance with procedures approved by the University of Pennsylvania Institutional Review Board.

### MRI acquisition

Participants were scanned on a Siemens 3.0 T Trio scanner. We acquired high-resolution T1-weighted structural images using an MPRAGE protocol (TR = 1620 ms, TE = 3.9 ms, flip angle = 15°, 1 mm slice thickness, 192 x 256 matrix, 160 slices, resolution = 0.9766 x 0.9766 x 1 mm). There were 3 functional scanning runs using gradient echo EPI sequences (32 slices in descending order of 3 mm thickness, a between slice gap of 0.75 mm, a resolution of 3 x 3 x 3 mm, a matrix size of 64 x 64, a flip angle of 78?, a TR of 2 s, and a TE of 30 ms). Each functional run lasted approximately 15 minutes.

### Stimuli

Stimuli were colored objects presented on a phase-scrambled background. Three categories of objects (apples, leaves, and roses) were presented in five colors (red, pink, yellow, blue, and green). There was also a warped, non-object condition that was presented in the same five colors. Examples of stimuli from each condition are displayed in Figure 1. To create the stimuli, high-resolution images of natural objects were edited in Adobe Photoshop. The background was removed, leaving an object in isolation. The portion of the object containing the relevant color property was manually segmented and placed into a separate layer, where we were able to modify its color independent of the other object features (e.g., for an apple image, the body of the apple was segmented and its color was modified without altering the stem or the leaves). This segmented portion of the object was first set to grayscale. We then created colored versions of each object by modifying the RGB color settings for this grayscale segmentation to red (RGB: 121 18 21), pink (RGB: 222 103 147), yellow (RGB: 187 174 30), blue (RGB: 0 67 166), and green (RGB: 0 171 0). Each object appeared in all five colors, ensuring that shape information was the same across all color conditions for a given object category. We repeated this procedure for 27 unique images within each object category (i.e., 27 apples, 27 leaves, and 27 roses). The same procedure and color settings were used for all objects. We also created mirror-flipped versions of the colored objects, resulting in 54 unique stimuli for each color-object condition (producing a total of 810 unique object stimuli). We created non-object images by applying a diffeomorphic warping procedure to the object stimuli described above. This procedure involves a smooth and continuous image transformation applied iteratively (40 iterations were used), and preserves low-level perceptual properties of the stimuli while making them unrecognizable as real-world objects [31]. All objects and non-object stimuli were centrally placed over a grayscale phase-scrambled background (the same background was used for all images).

### Stimulus presentation

We presented 810 unique object images to participants while collecting fMRI data from 15 categories of color-and-object combinations (Figure 1). Stimuli were presented in an event-related design using a continuous carry-over sequence within each run [32]. In each of the three runs, subjects viewed 270 unique object images (18 unique examples x 15 color-object conditions), as well as 36 unique non-object images. There were also 18 null events (5 s) in each run (null events were treated as an additional condition in the continuous carryover design) [32]. Each stimulus was presented on the screen for 1 s with an inter-stimulus interval of 1.5 s. On each trial subjects indicated by button press whether the image was an object or a non-object foil (Supplementary Figure S1). Therefore, subjects responded yes to all object images, regardless of typicality or category, and no to the non-object foils. Task accuracy was high. For object images the mean accuracy was 99.9% (SD = 0.1%), and for non-object images the mean accuracy was 96.4% (SD = 3.7%).

### Regions of interest

We defined a series of bilateral regions of interest (ROI) along the ventral visual pathway. These included ROIs for early visual cortex (EVC), V4, lateral occipital complex (LOC), inferior temporal gyrus (ITG), fusiform gyrus (FG), and perirhinal cortex (PRc). The ITG and FG ROIs were taken from the AAL atlas [33]. The EVC and LOC ROIs were taken from probabilistically defined parcels of functional localizer contrasts from a large number of subjects in a separate experiment (shared by the Epstein lab and described here [34]). These parcels were created through an automated procedure that identifies clusters of common activation across individuals for a series of functional ROI contrasts [35]. The LOC parcel was created from a contrast of objects > scrambled images, and the EVC parcel was created from a contrast of scrambled images > objects. We used the entire parcels for both EVC and LOC, and we did not apply any further voxel-selection procedures to these ROIs. Our V4 ROI was creating by placing spheres with a 6-mm radius around MNI coordinates that were reported in a classic study of color-perceptual processing [15], and which have previously been used to define ROIs for color processing [36]. The perirhinal cortex ROI was taken from a probabilistic map of anatomic segmentations [37] and was threshold to include voxels with at least 30% overlap across subjects.

### fMRI preprocessing and modeling

The fMRI data were processed and modeled using SPM8 (Wellcome Trust Centre for Neuroimaging, London, UK) and MATLAB (R2014a Mathworks; The MathWorks, Natick, MA). For each participant, all functional images were realigned to the first image [38] and co-registered to the structural image [39]. The images were spatially smoothed using a 3 mm FWHM isotropic Gaussian kernel. We modeled voxel responses to all conditions in each run in a single general linear model. Low-frequency drifts were removed using a high-pass filter with a cutoff period of 128 sec, and auto-correlations were modeled with a first-order autoregressive model. The parameter estimates for each condition were then averaged across runs. The resulting images were whole-brain maps of the voxel responses to each condition, which we then normalized to standard Montreal Neurological Institute space using a unified segmentation approach [40]. We used these maps to characterize the multivoxel information content in a series of ROIs through representational similarity analysis [41].

### Representational similarity analysis

We used representational similarity analysis (RSA) [41] to characterize the information encoded in the population responses of ROIs throughout the ventral visual pathway. For each ROI we constructed neural representational dissimilarity matrices (RDM) that represented all pairwise comparisons of conditions within each object category. The responses within each voxel were first z-scored across conditions, and we then computed neural dissimilarity as one minus the Pearson correlation coefficient between the multivoxel activation patterns for each condition. We also constructed dissimilarity matrices that represented the distances between conditions based on two models of representational content. The specifics of these two models are discussed below. We tested how well each model accounted for the representational structure in an ROI by calculating the Spearman correlation between the model and the neural dissimilarity matrices. The significance of each model was assessed using random-effects t-tests of the RSA correlations across subjects.

We examined two key models to test for the coding of a perceptual color space and a semantic color space. The perceptual model was created from subjective evaluations of color similarity collected in a norming study (described below). This model reflects the perceptual similarity of the colors independent of the object categories, and it was thus the same for each object category (e.g., red is more similar to pink than to green; Figure 2A). We converted these data into a dissimilarity matrix by taking the negative of the pairwise similarity values. The semantic model represents the dissimilarities between colors within each object category (e.g., red apple is more similar to green apple than to pink apple). Because the semantic statistics were unique to each object category, these models differed across categories (Figure 2B). Model fits were computed for each category separately, and we calculated the mean fit across categories. An important strength of this design is that all dissimilarity measurements reflect comparisons within an object category (i.e., apples, leaves, and roses), which means that the stimuli in each comparison contain the same shape information and only differ on color. This completely controls for shape information in both the perceptual and the semantic color models.

### Perceptual color model

We constructed a model of perceptual color similarity using subjective evaluations collected in a separate norming survey (N=18). This model captures color similarity independent of object categories. We presented the subjects with colored squares using the same RGB values used for the colored object images. Subjects judged the color similarity of the color swatches in a forced-choice two- alternative task with a reference swatch shown at the top and two choice swatches shown below. In an example trial, a subject might be shown a pink square at the top of the screen and asked to judge which of the two squares on the bottom, a red square or a blue square, is more similar. We constructed all possible pairings of reference and choice swatches (30 triads total), resulting in an equal number of judgments for all pairwise comparisons of colors. We used these data to construct a similarity matrix. For each pairwise comparison in this matrix, we counted the number of times that subjects reported those two colors as similar across all trials of the similarity judgment task. In other words, we filled the cells of this matrix with frequency counts of similarity pairings. We then converted this into a dissimilarity matrix by taking the negative of the similarity values. The resulting matrix captures color relationships that are closely matched to the perceptual space of a color wheel, as can be seen in the two-dimensional embedding in Figure 2A (all visualizations of two-dimensional embeddings were generated using t- distributed stochastic neighbor embedding [42]).

### Semantic color model

We constructed a model of semantic color similarity based on the feature co-occurrence frequencies for the colors and object categories. This model reflects color similarity relationships that are unique to each object category (e.g., green apples are more similar to red apples than to blue apples based on how frequently apples occur in these colors). We used a metric of co-occurrence frequency that captures the statistics of how people talk about object colors in written text [43]. This metric is derived from billions of words of text, and we reasoned that it would be strongly tied to how people think about and interpret these objects in the natural environment. We measured co-occurrence frequencies using Google ngram, a large corpus of English-language books [44]. Specifically, we quantified the directional co-occurrence frequencies of the color and object terms using both the singular and plural forms of the object terms (e.g., “red apple” and “red apples”) from 2008 (the most recent available data). We used log-transformed values of the co-occurrence statistics. To verify that these co-occurrence statistics related to the semantic interpretation of the objects, we asked the participants from the fMRI experiment to complete a series of subjective typicality ratings for the object images at the end of the study. The ratings were made on a 1-to-7 scale of highly atypical to highly typical. We were specifically interested in assessing whether subjects’ intuitions about color typicality related to the co-occurrence statistics of the object and color terms (e.g., that the high co-occurrence of “red apple” in text corresponded to subjective ratings that this color and object combination was highly typical). Indeed, there was a strong correlation between the co-occurrence statistics and mean subjective ratings of typicality across all objects (r = 0.71, p = 0.001). These co-occurrence statistics were then used to construct a model dissimilarity matrix of the semantic color space for each object category. We calculated the relative difference in co-occurrence for all pairwise comparisons of objects within a category (i.e., the absolute difference divided by the sum of the co-occurrence statistics). These dissimilarity matrices capture a model in which highly typical colors for an object category are close together in representational space, as can be seen in the two-dimensional embedding in Figure 2B [42].

## Supplemental Information

**Supplementary Figure S1.**
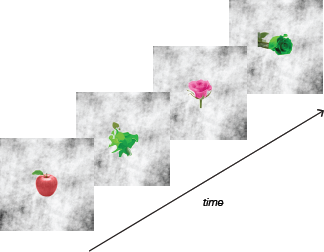
Visual-object behavioral task. Related to Figures 1 and 2. On each trial, participants viewed a single image on a phase-scrambled background and had to decide whether it was an object image or a warped non-object image.

**Supplementary Figure S2.**
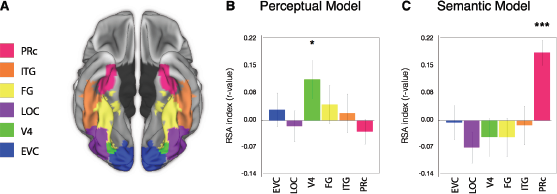
Representations of perceptual-color similarity and semantic-color similarity along the ventral visual stream. Related to Figure 3. To test for possible effects in other regions of the ventral stream, we performed the same analyses in a series of regions along ventral occipital-temporal cortex. (A) Color-coded regions of interest (B) Results for the perceptual-color model. The only region to show an effect for this model was V4 (p = 0.02). EVC: t(15) = 0.69, p = 0.50; LOC: t(15) = 0.33, p = 0.75; ITG: t(15) = 0.46, p = 0.65; FG: t(15) = 1.14, p = 0.27. (C) Results for the semantic model. The only region to show an effect for the semantic model was perirhinal cortex (p < 0.001). EVC: t(15) = 0.11, p = 0.91; LOC: t(15) = 1.71, p = 0.11; ITG: t(15) = 0.25, p = 0.81; FG: t(15) = 0.86, p = 0.40). Bar plots depict means ±SE. EVC = early visual cortex, LOC = lateral occipital complex, FG = fusiform gyrus, ITG = inferior temporal gyrus, PRc = perirhinal cortex. *p<0.05, ***p<0.001.

**Supplementary Figure S3.**
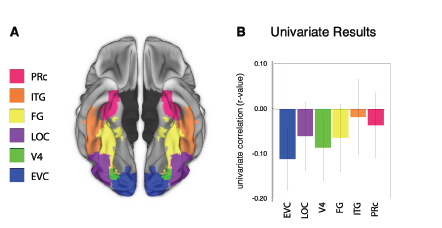
Summary of results from univariate analysis in all regions. Related to Figure 3. This analysis examined possible correlations between the typicality of color-and-category combinations and univariate activation, which might reflect a coarse familiarity signal within each object category. We calculated the correlation between the mean activity of each ROI and the log n-gram frequency of the color and object combinations. The correlation was calculated across all conditions after mean-centering the data within each object category (in order partial out overall differences across object categories and specifically examine the within-category effects of color typicality). Significance was assessed using random-effects t-tests of the univariate correlations across subjects. (A) Color-coded regions of interest. (B) We found no reliable correlation between color frequency and univariate signal in either PRc or V4 or in any of the other ventral visual regions of interest. (EVC (t(15) = 1.61, p = 0.13); LOC (t(15) = 0.80, p = 0.44); V4: t(15) = 1.18, p = 0.26; ITG (t(15) = 0.22, p = 0.83); FG (t(15) = 0.86, p = 0.40) PRc: t(15) = 0.50, p = 0.62). Bar plots depict means ±SE. EVC = early visual cortex, LOC = lateral occipital complex, FG = fusiform gyrus, ITG = inferior temporal gyrus, PRc = perirhinal cortex.

